# The cohesin subunit RAD21.2 functions as a recombination silencer of ribosomal DNA arrays

**DOI:** 10.1101/2022.06.20.496767

**Authors:** Viola Kuttig, Jason Sims, Yuki Hamamura, Shinichiro Komaki, Maren Köhler, Sara Christina Stolze, Joke De Jaeger-Braet, Hasibe Tuncay Elbasy, Hirofumi Nakagami, Antonio Virgilio Failla, Peter Schlögelhofer, Arp Schnittger

**Affiliations:** University of Hamburg, Department of Developmental Biology, Ohnhorststr. 18, D-22609 Hamburg, Germany; Department of Chromosome Biology, Max Perutz Labs, Vienna Biocenter, University of Vienna, Vienna, Austria; Nara Institute of Science and Technology, Graduate School of Biological Sciences, 8916-5 Takayama, Ikoma, Nara 630-0192, Japan; Max-Planck-Institute for Plant Breeding Research, 50829 Cologne, Germany; Light microscopy facility, University medical center Hamburg Eppendorf, Martini Straße 52, 22046 Hamburg

## Abstract

In many species, including Arabidopsis, heterochromatin often comprises repetitive DNA elements, such as arrays of ribosomal DNA (rDNA). Repetitive regions pose a risk in meiosis since recombination between them can lead to gross genomic rearrangements. However, meiotic recombination at rDNA arrays and other heterochromatic repeat regions is blocked by not well understood mechanisms. Here, we have identified RAD21.2, an α-kleisin subunit of cohesin, as a repressor of meiotic recombination at the rDNA regions in Arabidopsis. We show that RAD21.2 co-localizes with heterochromatic factors and is specifically enriched at rDNA repeats, which are devoid of the meiosis specific α-kleisin REC8, needed for recombination. Knocking down RAD21.2, we find that REC8 moves into the nucleolus organizing regions (NORs), where we see an increase of RAD51 recombinase foci numbers. Concomitantly, we find extensive rearrangements of the NORs and the offspring of these plants have large variation in rDNA copy numbers demonstrating that RAD21.2 is necessary for transgenerational genome stability.

**One-Sentence Summary:** The cohesin component RAD21.2 represses meiotic recombination and by that contributes to genome stability over generations.

## Main Text

According to a broadly accepted model, cohesin complexes embrace the two sister chromatids of each replicated chromosome with their ring-like structure^1^. Cohesins are essential for two central aspects of meiosis^1^: First, they mediate the ordered segregation of homologous chromosomes (homologs) in meiosis I and sister chromatids in meiosis II to yield balanced gametes with half of the DNA content of the meiotic mother cell. Second, cohesins are an integral part of the meiotic chromosome axis and are therefore crucial for meiotic recombination and genetic diversity of the offspring.

The core cohesin complex is composed of four subunits: SMC1 and SMC3, two ATPases that belong to the family of structural maintenance of chromosomes (SMC) proteins, the HEAT-repeat domain protein SCC3/SA and an α-kleisin component^1^. To accommodate meiosis-specific functions, the mitotic α-kleisin RAD21 is usually replaced by REC8 in meiosis^2–4^. Yet, REC8 is not the only meiosis-specific α-kleisin and additional α-kleisins have been identified to be relevant for meiosis in *C. elegans* and mammals^5–9^. Up to now, these α-kleisin components have been found to work together with and/or take over specific functions of REC8 in mediating cohesion and recombination. However, additional α-kleisin components are prevalent in other eukaryotes and their role is not very well understood.

In addition to REC8 (also called SYN1 or DIF1), the model plant *Arabidopsis thaliana* encodes three predicted α-kleisins named RAD21.1 (SYN2), RAD21.2 (SYN3), and RAD21.3 (SYN4). Currently, the specific function of these three *RAD21* genes is poorly understood. *RAD21.1* and *RAD21.3* have been implicated in the DNA damage response. However, no severe mutant phenotype could be detected for single and double mutants^10^. In contrast, *RAD21.2* is an essential gene and homozygous mutants could not be recovered^10, 11^. Heterozygous mutants in *RAD21.2* show fertility defects and the protein seems to be required for both pollen and embryo sac development. In addition, both RNAi-mediated silencing of *RAD21.2* and overexpressing a C-terminally tagged RAD21.2 resulted in early meiotic defects, which include loss of synapsis and reduced loading of the transverse element ZYP1^12, 13^. Using an antibody, RAD21.2 was reported to localize to the nucleolus of meiocytes. Together with the additionally observed changes in protein accumulation patterns, these findings question whether RAD21.2 acts as a *bona fide* cohesin component^12^.

To determine the expression pattern of RAD21.1 and RAD21.3 and to revisit the accumulation of RAD21.2 in meiosis, we generated genomic reporter constructs in which GFP was inserted at the C-termini. To facilitate a co-localization analysis with REC8, we exchanged GFP with RFP in a previously published functional reporter for REC8 (*PRO_REC8_:REC8:GFP*)^14^. We found that all three RAD21 genes are expressed in somatic tissues where REC8, as expected, is absent (Fig. S1AI-II and BI-II, Fig. S2AI-II and Movie S1). In addition, all three RAD21 proteins accumulate in the progenitor cells of meiocytes and in the tissue surrounding the meiocytes (Fig. S1AIII, BIII and Fig. S2B). RAD21.1:GFP and RAD21.3:GFP were not detected in meiocytes, where instead REC8 is expressed (Fig. S1AIII and BIII). In contrast, RAD21.2:GFP is present in meiosis I and decorates meiotic chromosomes (Fig. S2B). A similar chromosomal localization pattern was also found for an N-terminal fusion to RAD21.2, i.e. *PRO_RAD21.2_:GFP:RAD21.2* (Fig. S2B). Importantly, the N-terminally tagged protein fully complements the growth and fertility defects observed in *rad21.2* mutants (Fig. S3A). The localization of RAD21.2 is consistent with its potential function as a cohesin subunit^12^.

To address whether RAD21.2 could act as a cohesin subunit, we first revealed that it was binding in yeast two-hybrid assays to the core cohesin components SMC1 and SCC3 (Fig. S3B). Next, we tested whether this interaction can also be found *in vivo*. To this end, we immunoprecipitated GFP-fused RAD21.2 from transgenic seedlings and determined proteins interacting *in vivo* by mass spectrometry analysis. As a control, we used plants expressing unfused GFP. SMC1 and SMC3 were identified as the top 2 significantly enriched proteins pulled down with RAD21.2 (Fig. S4A, Table S1, SMC1: p=0.003, SMC3: p=0.001). Thus, we conclude that RAD21.2 is part of a cohesin complex *in vivo*.

Next, we aimed at a detailed localization analysis of RAD21.2 throughout meiosis. However, the fluorescent signals of RAD21.2:GFP and GFP:RAD21.2 are not sufficiently intense when expressed under the control of the endogenous *RAD21.2* promoter to conduct live cell imaging experiments. Therefore, we generated a further *RAD21.2* reporter construct using the *ASK1* promoter to drive *GFP:RAD21.2* (*PRO_ASK1_:GFP:RAD21.2*). The construct could fully complement the deficiencies of homozygous *rad21.2* mutant lines (Fig. S3A). Importantly, the localization pattern of the GFP:RAD21.2 fusion protein expressed from the *ASK1* promoter could readily be assessed and showed qualitatively the same chromosome association as GFP:RAD21.2 expressed from its endogenous promoter (Fig. 1A, Movies S2 and S3). For reasons of simplicity the *PRO_ASK1_:GFP:RAD21.*2 plant line is termed a*GFP:RAD21.2* below.

**Fig. 1.**
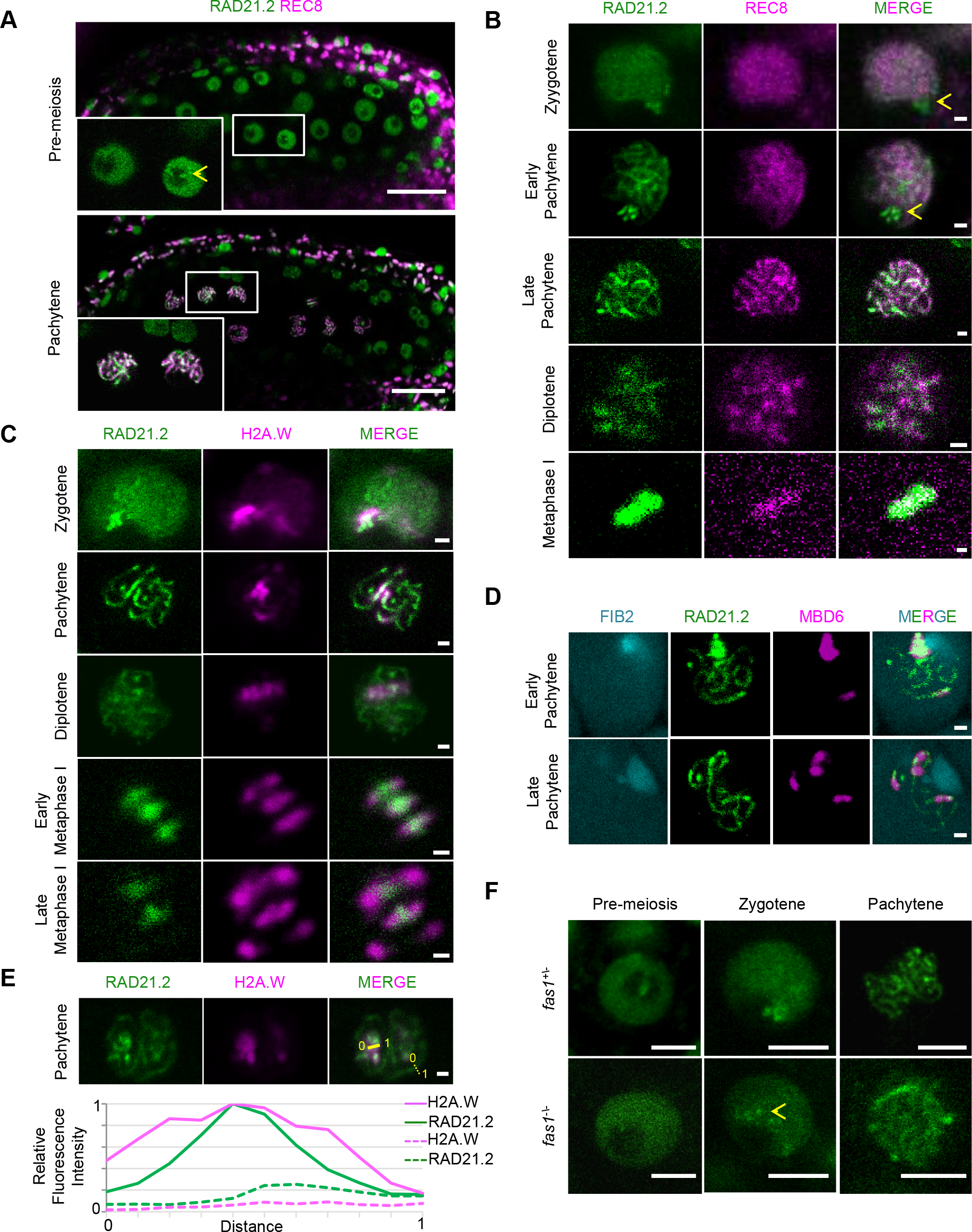
Meiotic localization pattern of RAD21.2: **A,** Confocal laser scanning micrographs of Arabidopsis anthers expressing PRO_REC8_:REC8:RFP (magenta) together with PRO_ASK1_:GFP:RAD21.2 (green). Upper row, RAD21.2 but not REC8 is present in all cells prior to meiosis (close-up highlighted in box). Note the thread like structure decorated by RAD21.2 reaching into the nucleolus (arrowhead), see text for details. Lower row, next to REC8, only RAD21.2 out of the three RAD21 proteins accumulates in meiosis (here pachytene stage) and decorates chromosomes. Scale bar: 20 µm. **B**, Confocal laser scanning micrographs of male meiocytes revealing distinct and largely not overlapping localization patterns of PRO_ASK1_:GFP:RAD21.2 (green) and PRO_REC8_:REC8:RFP (magenta). Notably, RAD21.2 is enriched at nucleolar chromatin (arrowhead) in early prophase I. Scale bar: 1 µm. **C**, Confocal laser scanning micrographs of male meiocytes showing that PRO_H2A.W.6_:H2A.W.6:RFP (magenta) largely co-localizes with a sub-fraction of the PRO_ASK1_GFP:RAD21.2-marked chromatin (green), for instance in the perinucleolar region (arrowheads). Scale bar: 1 µm. **D**, Confocal laser scanning micrographs of meiocytes expressing the nucleolus reporter PROFIB2:FIB2:mTurquoise (cyan), PRO_ASK1_:GFP:RAD21.2 (green) and PRO_HTR5_:MBD6:RFP (magenta) showing highly methylated DNA regions, which are decorated by RAD21.2, clustered in the proximity of the nucleolus at early pachytene (upper row) and a distributed pattern during the dissolution of the nucleolus in late pachytene (lower row). Scale bar: 1 µm. **E**, Quantification of the overlap between H2A.W.6 and RAD21.2 accumulation patterns seen in **C**. Upper row: Yellow lines indicate a region in the perinuclear region (solid line) and a region distant from the nucleolus (dashed line) used for quantification. Lower row: Profile plot of the relative fluorescence intensities of RAD21.2 (green lines) and H2A.W (magenta lines) in the nucleolar area (solid lines) and outside of the nucleolus (dashed lines). The fluorescence intensities were normalized to the highest fluorescent value. Scale bar: 1 µm. **F**, Confocal laser scanning micrograph of an anther expressing *PRO_ASK1_:GFP:RAD21.2* in *fas1^+/-^* mutants show the typical chromosomal localization pattern of GFP:RAD21.2 (green) from pre-meiosis to pachytene. A compromised localization in *fas1* mutants was observed starting from zygotene onwards. Scale bar: 5 µm.

We combined the a*GFP:RAD21.2* line with our *REC8:RFP* reporter and found that RAD21.2, in contrast to the even distribution of REC8 along chromosomes, largely accumulates on chromosomes at the border of the nucleolus from leptotene through zygotene until early pachytene (see arrowhead Fig. 1B). A similar localization, albeit much weaker, is also seen for the *PRO_RAD21.2_:RAD21.2:GFP* (Fig. S2B). Towards late pachytene, the cluster of RAD21.2 cannot be detected and becomes more diffusely distributed in the nucleus. These results are consistent with the notion that the regions enriched in RAD21.2 are associated with the nucleolus, where REC8 is not present (Fig. 1B). The nucleolus association was further supported by analyzing a reporter line containing a *GFP:RAD21.2* together with a C-terminally TFP-tagged *FIBRILLARIN 2* gene (*FIB2*) expressed under its endogenous promoter as a nucleolus marker (*PRO_FIB2_:FIB2:TFP*)^15^ (Fig. 1D and S4B).

During metaphase I, the local fluorescence intensity of aGFP:RAD21.2 increases likely due to the condensation of the chromosomes (Fig. 1B, C and Movie S3). At the onset of anaphase I, the aGFP:RAD21.2 signal completely disappears consistent with the cleavage of RAD21.2 by separase. We never observed re-appearance of aGFP:RAD21.2 fluorescence during meiosis after anaphase I. However, aGFP:RAD21.2 can be detected again once meiosis is completed in the developing microspores (Fig. S4C).

To confirm the localization of RAD21.2, we performed Lipsol spreads of wild-type meiocytes staining for RAD21.2 (newly generated antibody – this study) and for REC8. The results show a very similar localization pattern of RAD21.2 to the aGFP:RAD21 proving that aGFP:RAD21 is not only functional but shows an accumulation pattern similar to that of the endogenous RAD21.2. Furthermore, we also performed super-resolution stimulated emission depletion (STED) microscopy of RAD21.2 and REC8 revealing the structure of the RAD21.2 cluster which is located at regions completely depleted from REC8. This further supports our findings that the region to which RAD21.2 is binding localizes to the NORs (Fig. S4D).

Next, we asked whether the correct localization of RAD21.2 would be dependent on the removal of REC8 from the nucleolus-associated regions. REC8 in Arabidopsis is subject to the prophase pathway of cohesin removal mediated by the AAA+ ATPase WAPL^18, 19^. We tested the localization of RAD21.2 in the absence of REC8 removal. However, we did not see any alteration of the aGFP:RAD21.2 signal intensity and distribution in *wapl1 wapl2* mutants indicating that the localization of RAD21.2 does not depend on the removal of REC8 and that RAD21.2 itself, in contrast to REC8, is not subject to the prophase cohesin removal pathway (Movie S4 and Fig. S5A). Conversely, we did not find any obvious differences in the distribution of aGFP:RAD21.2 when expressed in *rec8* mutants (which remained not fertile, Fig. S5B-C) compared to the wildtype (Fig. S5D).

Since the NOR regions associated with the border of the nucleolus are comprised of heterochromatin, we next tested to what extent RAD21.2 is co-localizing with GC methylation as a hallmark of heterochromatin (visualized by *PRO_HTR5_:MBD6:RFP*)^16^, and with the histone variant H2A.W, which is specifically incorporated in heterochromatic domains (*PRO_H2A.W.6_:H2A.W.6:RFP*)^17^. Our results show that both heterochromatin markers strongly overlap with RAD21.2 (Fig. 1CD-E). Analyzing RAD21.2 together with H2A.W.6 in somatic cells (Fig. S2B) also shows a strong co-appearance of RAD21.2 and heterochromatic domains. This hypothesis was further corroborated by the observation that RAD21.2 strongly overlapped with late replicating DNA regions known to represent heterochromatin^20^, as visualized by the co-localization of RAD21.2 with large speckles of PCNA:RFP formed in late S-phase (Fig. S2D). Taken together, we conclude that the loading of RAD21.2 in meiocytes correlates with a heterochromatin environment.

To explore if RAD21.2 loading depends on a heterochromatic environment, we analyzed the localization of RAD21.2 in plants mutant for the gene *DEFICIENT IN DNA METHYLATION 1* (*DDM1*), which encodes a chromatin-remodeling protein that is required for the maintenance of heterochromatin^21^ and genome stability^22^. Recently, it has been reported that DDM1 binds to H2A.W and mediates its deposition to heterochromatin^23^. However, an obvious change of the localization of RAD21.2 in *ddm1* mutants was not observed (Fig. S6A). Next, we analyzed the localization of RAD21.2 in *nucleolin 2* (*nuc2*) mutants since the large RAD21.2 cluster is nucleolus-associated. NUC2 is required for chromatin organization of silent 45S rDNA, and its loss leads to major changes in trans-generational stability of the 45S rDNA^24^. No obvious changes could be detected in the localization of RAD21.2 when compared to wild type (Fig. S6A). In contrast, the RAD21.2 signal was reduced in mutants of the *FASCIATA 1* (*FAS1*) gene from pre-meiosis throughout prophase I (Fig. 1F). *FAS1* encodes for a subunit of the CHROMATIN ASSEMBLY FACTOR (CAF) which is required for nucleosome assembly and maintenance of heterochromatin particularly the 45S rDNA repeats^25, 26^. Notably, we never saw the typical RAD21.2 accumulation close to the nucleolus at zygotene/early pachytene (Fig. S6A, arrowhead). This could also reflect the lower copy number of the 45S rDNA repeat present in *fas1* mutants^25, 26^. Thus, proper accumulation of RAD21.2 appears to depend on the correct nucleosome assembly in heterochromatic regions and on a wild-type copy number of the rDNA repeats.

Since the aGFP:RAD21.2 shows a clustered localization in the premises of the nucleolus, we investigated whether RAD21.2 would co-localize with the 45S rDNA region. To this end, we performed an Immuno-FISH experiment on Lipsol spreads of PMCs targeting RAD21.2, ASY1 and the 45S rDNA. The results showed that the RAD21.2 cluster colocalizes with the 45S rDNA indicating that there is an overabundance of this specific cohesion subtype at the rDNA region (Fig. S6B). Furthermore, RAD21.2 overlaps with only a part of the 45S rDNA reinforcing the idea that it is associated with the heterochromatic regions of the NORs.

To probe the functional relevance of the RAD21.2 occupation at the nucleolar associated domains, we next generated an RNAi construct against *RAD21.2* since loss of RAD21.2 results in gametophytic lethality^11^, precluding an easy assessment of the consequences of altered RAD21.2 abundance in meiosis. We recovered two independent transgenic lines with 25-35 percent lower *RAD21.2* expression levels that exhibit no obvious vegetative growth defects (Fig. S7A, B) but showed a reduction in silique length (Fig. S7C). The knock-down lines revealed a reduction of about 30% in pollen viability and a seed abortion level of around 45% (Fig. S7D, E). To address whether the defect does arise from chromosomal translocations generated by the insertion of the RNAi construct, we crossed the RAD21.2 RNAi #1 with the wildtype and performed spreads on pollen mother cells. We could not detect any univalents or mispaired chromosomes. These results, together with the similarity between the two independent RNAi lines confirm that the effects we detect are genuinely due to the knock-down of *RAD21.2* (Fig. S7F).

To assess whether the reduced fertility in the RAD21.2 RNAi lines is due to defects occurring during meiosis, we performed spreads on pollen mother cells (PMCs) (Fig. 2A and S8A). In wild-type plants, 5 separated bivalents are visible at diakinesis and metaphase I. In contrast, *RAD21.2 RNAi* plants show severe chromosomal defects. Entanglements and connections between non-homologous chromosomes could be observed. During metaphase I, most cells of wild-type plants formed five distinct bivalents (cells with chromosomes entanglements: 15%; n=130), whereas *RAD21.2 RNAi* plants showed at least two connected chromosome pairs per cell with a stretched morphology at a high frequency (cells with chromosomes entanglements: 72%; n=130) (Fig. 2B). We also observed defects in the second meiotic division with 7% of the meiocytes in the *RAD21.2 RNAi* plants showing unbalanced chromosome numbers at metaphase II (n=28) (Fig. 2A), while no incident of unbalanced chromosomes was found in the wild-type (n=24).

**Fig. 2.**
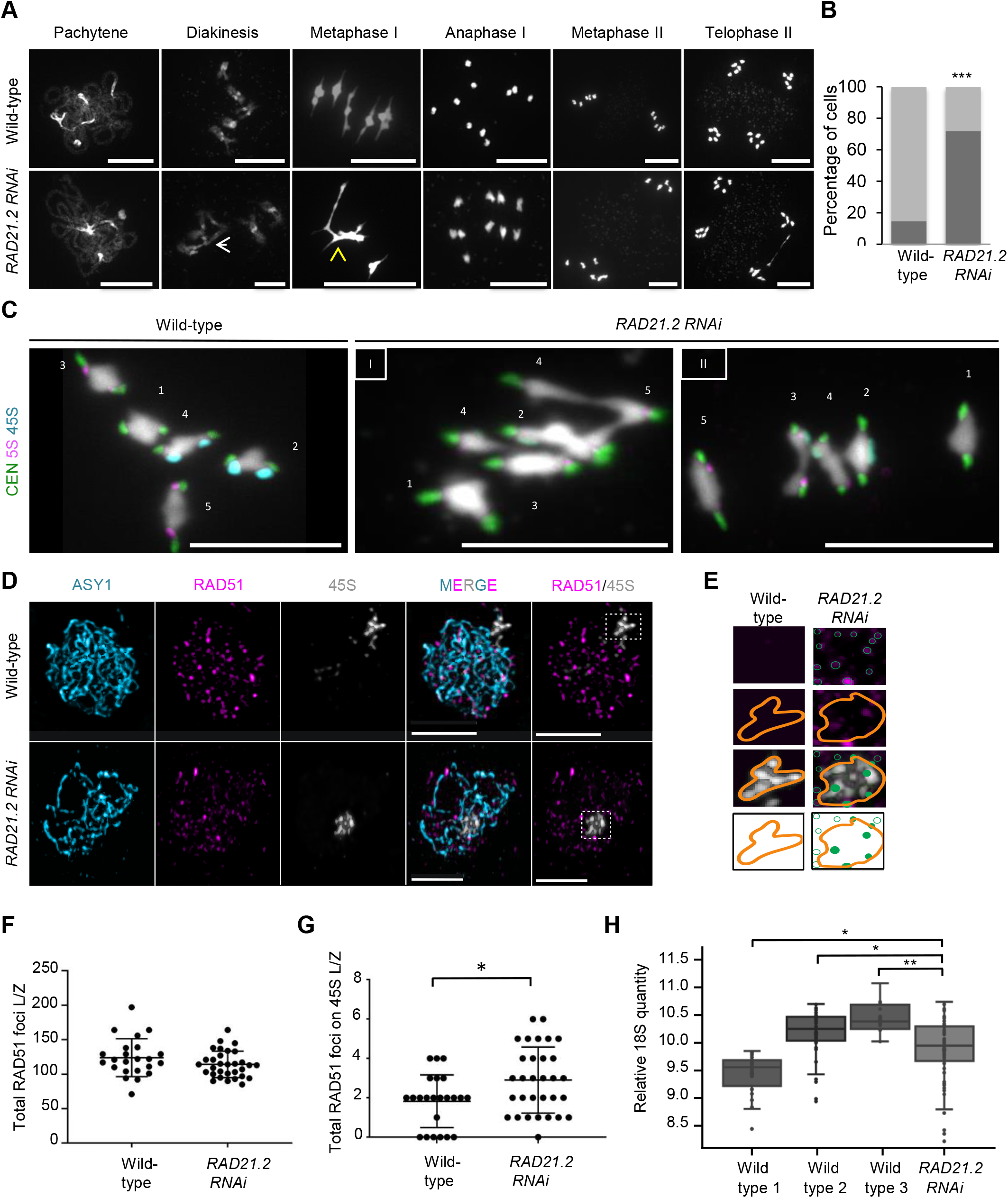
Knocking down of *RAD21.2* results in recombination defects: **A,** Chromosome spread analysis of pollen mother cells in wild-type (upper row) in comparison to *RAD21.2 RNAi* plants (lower row), which often have chromosome entanglements seen in diakinesis and metaphase I (arrowhead). Scale bar: 10 µm. **B**, Graph depicting the percentage of cells with (dark grey) and without (light grey) chromosome entanglements in metaphase I of the wild-type (15%; n=130) and *RAD21.2 RNAi* (72%; n=130; p-value=8.23E-27, Student’s *t*-test). **C**, FISH analysis of metaphase I cells of pollen mother cells from the wild-type and *RAD21.2 RNAi* plants. Probes against 45S rDNA (cyan), 5S rDNA (red) and CEN (green) loci were used to identify chromosomes; DNA was visualized by DAPI (grey). See text for details. Scale bar: 10 µm **D**, Immuno-FISH analysis of wild-type and *RAD21.2 RNAi* pollen mother cells at zygotene. The axis has been stained with anti-ASY1 (cyan) for staging and the DNA repair sites are highlighted by anti-RAD51 (magenta). The 45S rDNA has been visualized with a specific FISH probe (white). Scale bar: 5 µm. **E**, Related to **D**, RAD51 foci were counted in the NOR region, marked by the orange line. **F**, The total number of RAD51 foci at leptotene/zygotene stage in wild-type versus *RAD21.2 RNAi* plants is not significantly different. **G**, The number of RAD51 foci counted on the 45S region at leptotene/zygotene is significantly larger in *RAD21.2 RNAi* plants than in the wild-type. **H**, Box plot depicting the 18S gene copy number in the offspring of 3 wild-type plants (Wildtype 1: n=23, Wildtype 2: n=48, Wildtype 3: n=16) compared to the offspring of a *RAD21.2 RNAi* plant (n=78), which has a significant higher variance of the 18S copy number than the wild-type.

To address the nature of the chromosomal abnormalities at metaphase I, we performed fluorescence *in situ* hybridization (FISH) (Fig. 2C). We identified connections between non-homologous chromosomes, for instance between chromosomes 3, 4 and 5 (Fig. 2C) and between chromosomes 3 and 4 (Fig. S8CI/II). We also revealed more complex chromosomal rearrangements such as the two homologous chromosomes 4 connected to other non-homologous chromosomes (Fig. 2CI) and a genome rearrangement event involving the 45S rDNA region, which is translocated from chromosome 4 to chromosome 3 (Fig. 2CII). In addition, we observed fragments of the 45S rDNA after meiosis I (Fig. S8BIII, arrowhead). Furthermore, we identified connections between the centromeres of chromosomes 2 and 3, suggesting a general increase in genome instability involving repetitive DNA regions (Fig. S8CII).

Several of the above-described chromosomal rearrangements could be explained by recombination events between the rDNA regions in the absence of RAD21.2. To investigate this, we examined the localization of the recombinase RAD51 at the 45S rDNA region in leptotene/zygotene stages by immuno-FISH (Fig. 2D, E). While the total number of RAD51 foci in wild-type (123±27, n=23) and in *RAD21.2 RNAi* plants (114±18, n=31) are similar, the number of RAD51 foci at the 45S rDNA region increases from 1.4±1 foci in wild-type to 2.7±1.5 (p=0.0049) foci in the *RAD21.2 RNAi* meiocytes (Fig. 2F, G). This result supports the idea that RAD21.2 is needed to suppress meiotic recombination at the rDNA loci, especially given that the RAD21.2 RNAi plants, which could be recovered, only represent a moderate knock-down of RAD21.2.

As shown above and demonstrated in a previous study^27^, the meiotic α-kleisin REC8, which is key for meiotic recombination as a part of the chromosome axis, is mostly excluded from the 45S rDNA region. To determine the localization of REC8 in this region in the *RAD21.2 RNAi* meiocytes, we performed immuno-FISH using fixed meiocytes (Fig. 3A, B). Comparing the relative fluorescence intensity profile plots of REC8 at the 45S loci revealed an increase in abundance of REC8 signal at the 45S region in the *RAD21.2 RNAi* plants (n=15, p=0.032) compared to the wild-type (n=15) (Fig. 3C-E).

**Fig. 3.**
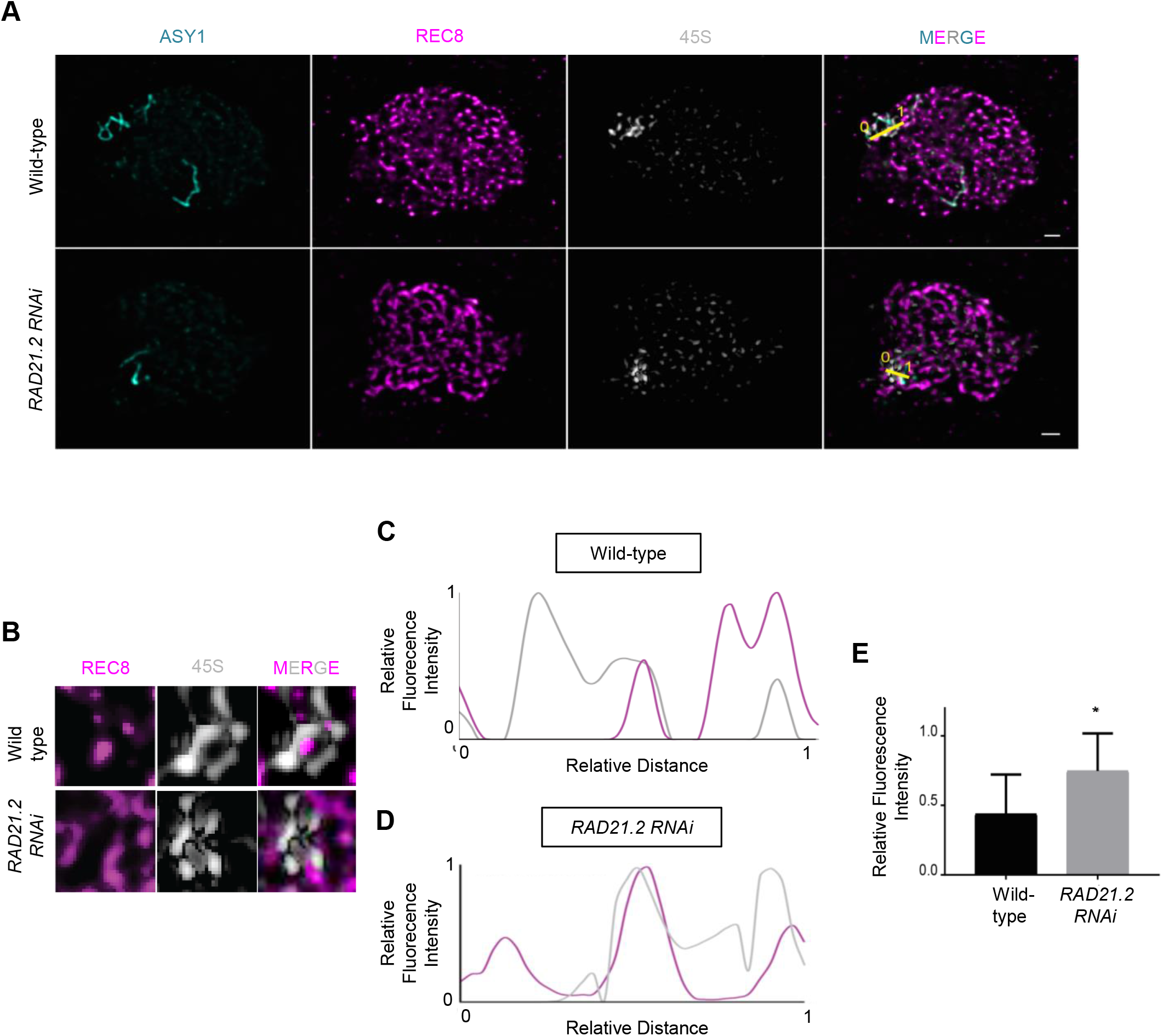
REC8 accumulates in the 45S rDNA region of meiocytes in *RAD21.2 RNAi* plants: **A,** Immuno-FISH analysis of wild-type and *RAD21.2 RNAi* pollen mother cells at pachytene. The axis has been stained with anti-ASY1 (cyan) and the meiosis-specific cohesin subunit with anti-REC8 (magenta). The 45S rDNA has been visualized with a specific FISH probe (white). The yellow lines define regions used to quantify the fluorescence intensities. Scale bar: 1 µm. **B**, Related to **A**, magnification of the 45S rDNA region for the wild-type and *RAD21.2 RNAi* plants with REC8 (magenta) and 45S (light grey). An increased REC8 localization to the 45S rDNA in *RAD21.2 RNAi* plants in comparison to the wild-type can be observed. **C** and **D**, Profile plots of the fluorescence intensities of REC8 (magenta) and 45S rDNA (light grey) for the wild-type (**C**) and *RAD21.2 RNAi* plants (**D**). The fluorescence intensity was normalized to the highest fluorescent value. **E**, The average relative fluorescence intensity of the REC8 signal taken at the maxima of the 45S rDNA is significantly higher in *RAD21.2 RNAi* plants (n=15) than in the wild-type (n=15) (p-value=0.032).

We reasoned that an increase in recombination in the rDNA region should also affect gene copy number through deletions and insertions. Indeed, the 18S rDNA gene copy number varied significantly more in the offspring of the *RAD21.2 RNAi* plant compared to the progeny of three different wild-type plants (Fisher’s F test, WT-1/*RNAi* p=0.018, WT-2/*RNAi* p=0.049 and WT-3/*RNAi* p=0.0045; Fig. 2H). To assess whether the 18S gene copy number variability is due to the rearrangements occurring in meiosis and not due to somatically occurring defects, we analyzed the 18S rDNA copy number in leaves of different sizes in fully grown plants. The results showed no difference in the 18S copy number indicating that the variations detected in the progeny of the RAD21.2 RNAi plants arises from meiotic defects and not somatically (Fig. S8D).

Taken together, we have revealed a novel function in meiosis for the so far poorly characterized α-kleisin component RAD21.2. RAD21.2 is specifically loaded on the heterochromatic domain of the rDNA and possibly other heterochromatic regions, where it prevents loading of REC8 and suppresses aberrant recombination events. The role of RAD21.2 as a REC8 repellent stands in striking contrast to the so far described functions of other meiotic α-kleisin components in other organisms, which co-operate with and/or substitute REC8 function^5–9^. It will now be interesting to analyze across different organisms to what degree other meiotic α-kleisins may function as anti-recombination factors in maintaining genome stability.

## Acknowledgments

We thank Eva R. Hoffmann (University of Copenhagen), Kostas Lampou (University of Hamburg), Lucas Lang (University of Hamburg), and Maren Heese (University of Hamburg) for critical reading and helpful comments to the manuscript. We thank Anne Harzen for support in the mass-spectrometry experiments. We are grateful to the Max-Planck-Gesellschaft (HN) and University of Hamburg (AS) for core funding. This work was supported through an Austrian Science Fund grant (# I 3685-B25) to P.S. and a DFG grant (SCHN 736/8-1) to A.S.

## Data and materials availability

Figs. 1 to 3

## Materials and Methods

### Plant material

In this study, the *Arabidopsis thaliana* accession Columbia (Col-0) was used as the wild-type reference. The used T-DNA insertion lines SALK_044851 (*rad21.1*), SALK_053140 (*rad21.2*), SALK_076116 (*rad21.3*), SAIL_807_B08 (*rec8*), SALK_076791 (*wapl1-1*), SALK_127445 (*wapl2*), GK_178D01 (*nuc2-2*), *ddm1-2* and SAIL_662_D10 (*fas1*) were obtained from the Nottingham Arabidopsis Stock Center (http://arabidopsis.info/). The *35S:AP1-GR ap1 cal* line was kindly provided by Frank Wellmer^28^. Genotypes were determined by PCR using primers listed in Supplementary Table S2. *PRO_REC8_:REC8:GFP*^14^ (*14*), *PRO_H2A.W.6_:H2A.W.6:RFP*^17^, *PRO_HTR5_:MBD6:GFP*^16^ and *PRO_RPS5_:RFP:TUA5*^29^ reporters were previously generated.

### Plant growth conditions

Seeds were surface-sterilized with chlorine gas and sown on 1% (w/v) agar plates containing half-strength Murashige and Skoog (MS) salts, 1% sucrose, pH 5.8. Antibiotics were added for seed selection when required. For stratification, plates were stored 2 days at 4°C in the dark, thereafter plates were transferred for 10 days to a growth chamber with long day conditions (16h of light; 21°C/ 8h of dark; 18°C and 60% humidity) for seed germination. Seedlings were transferred to soil and grown under long day conditions until seed production.

### Plasmid constructions and plant transformation

To create the *PRO_RAD21s_:RAD21s:GFP* and *PRO_RAD21.2_:GFP:RAD21.2* constructs, a fragment covering the genomic region of each gene together with an upstream region of the start codon of 2 Kb, 1 Kb and 2.5 Kb, respectively, along with 1 Kb downstream of the stop codon of each gene was amplified by PCR and cloned into pENTR2B by SLiCE. A restriction enzyme site (*Sma*I for *RAD21.1* and *RAD21.2*, and *Nae*I for *RAD21.3*) was inserted in front of the stop codon (C-terminal GFP fusion) or behind the start codon (N-terminal GFP fusion) of the RAD21s constructs. The resulting construct was linearized by the restriction enzyme digestion and was ligated to the *GFP* gene, followed by LR recombination reactions with the destination vector *pGWB501*.

For the exchange of the native RAD21.2 promoter with the ASK1 promoter (1 kb upstream of the start codon), the promoter sequence was amplified by PCR and cloned into the *pENTR2B PRO_RAD21.2_:GFP:RAD21.2* by SLiCE, followed by LR recombination reaction with the destination vector *pGWB501*.

To generate the *PRO_FIB2_:FIB2:mTurquoise,* the genomic FIB2 sequence and 1kb upstream of the start codon and 800 bp downstream of the stop codon was amplified by PCR and cloned into the *pENTR2B* vector by SLiCE. A *SmaI* restriction enzyme site was inserted in front of the stop codon. The resulting construct was linearized by *SmaI* digestion and was ligated to the *mTurquoise* gene. To generate the *RAD21.2 RNAi* construct, a 400 bp fragment of the *RAD21.2* CDS was amplified by PCR with attB flanking primers and cloned into the *pDONR221* vector by gateway BP reaction. The resulting construct was integrated into the *pK7GWIWG2* vector by gateway LR reaction. All primers used for plasmid construction are listed in Supplementary Table S2.

All constructs were transformed into *Arabidopsis thaliana* plants by floral dipping.

### Phenotypic evaluation

Peterson staining was used to analyze the pollen viability (Peterson et al., 2010). Three flower buds containing either dehiscent or non-dehiscent (for whole anther staining) pollen were collected and dipped in 25 µl Peterson staining solution (10% ethanol, 0.01% malachite green, 25% glycerol, 0.05% acid fuchsin, 0.005% orange G, 4% glacial acetic acid) for 15 s on a microscope slide that was covered by a coverslip. Slides were incubated at 80°C for 10 min (for pollen counting) or 30 min (for whole anther staining) and aborted and non-aborted pollen grains were observed using a light microscope. Seed sets were determined by quantifying viable and aborted seeds of mature siliques; 3 siliques per plant were analyzed.

### Cytogenetic analysis

The preparation of pollen mother cells DAPI spreads was performed as previously described^24^. Flower buds were fixed in 3:1 ethanol/ acetic acid (fixative) over night and washed once with fresh fixative solution followed and stored in 70% ethanol at 4°C. The flower buds were staged by size and washed once with ddH20 and once with 10mM citrate buffer. The digestion of flower buds was performed in 10 mM citrate buffer (0.5% w/v cellulose, 0.5% w/v pectolyase and 0.5% w/v cytohelicase) for 2.5 hours at 37 °C. For the chromosome spreading, single flower buds were transferred to a drop of 45% acetic acid on a glass slide and squashed with a bended needle for 1 min. The spreading was performed for 1 min on a 46°C hot plate. The slide was washed with fixative solution and dried for at least 2 hours. The chromosome spreads were stained by 18 µl of Vectashield Antifade Mounting medium with DAPI (vector laboratories) and sealed with a cover slip.

#### FISH

The DAPI slides selected for fluorescence *in situ* hybridization (FISH) were washed in 100% ethanol until the coverslips could be easily removed (5-10 min) and subsequently washed in 4T (4X SCC and 0.05% v/v Tween20) for at least 1 h in order to remove the mounting medium.

After washing the slides in 2X SCC for 10 min they were placed in pre-warmed 0.01 M HCl with 250 µl of 10 mg/ml Pepsin for 90 seconds at 37 °C. The slides were then washed in 2X SCC for 10 min at room temperature. 15 µl of 4% paraformaldehyde (PFA) were added onto the slides, covered with a strip of autoclave bag and placed for 10 min in the dark at RT. The slides were then washed with deionized water for 1 minute and dehydrated by passing through an alcohol series of 70, 90, 100 %, for 2 minutes each. Slides were left to air-dry for 30 min.

Meanwhile, the probe mix was prepared by diluting 1 µl of probe (2-3 µg of DNA) in a total of 20 µl of hybridization mix (10% dextran sulphate MW 50,000, 50% formamide in 2x SSC).

Only 50 pmols (final concentration) of the LNA probes were used per slide. The probe mix was denatured at 95 °C for 10 min and then placed on ice for 5 min. Afterwards, the probe mix was added to the slide, covered with a glass coverslip, sealed and placed on a hot plate for 4 min in the dark at 75 °C. Finally, the slides were placed in a humidity chamber over-night at 37 °C. After hybridization, the coverslips were carefully removed and the slides were treated with 50% formamide in 2X SCC for 5 min in the dark at 42 °C. The slides were then washed twice with 2X SCC for 5 min in the dark at room temperature. Finally, 15 µl of DAPI-Vectashield solution were added to the slide and sealed with a coverslip. Images were taken on a Zeiss Axioplan microscope (Carl Zeiss) equipped with a mono cool-view CCD camera. For all repetitive regions analyzed we used specific LNA probes see Table S2.

### RAD21.2 antibody generation

Polyclonal antibodies against peptides CET GPD NEP RDS NIA and CNW ETE SYR TEP STS T were generated in rat and affinity purified. The affinity purified RAD21.2 antibody was used in a dilution of 1:5 in blocking solution for immuno-histochemistry. The peptide synthesis, animal immunization and affinity purification were outsourced to Eurogentec.

### Immuno-FISH

Immuno-FISH was performed using the TACE method (*24*). Immunofluorescence (IF) antibodies were used as follows: anti-ASY1 raised in guinea pig 1:10,000, anti-RAD51 raised in rat 1:300, anti-RAD21.2 raised in rat 1:5 (affinity purified), anti-REC8 raised in rabbit 1:250, anti-guinea pig Alexa488 (Abcam #ab150185) 1:400, anti-rat Alexa568 (Abcam #ab175476) 1:400. 45 rDNA was detected by using an LNA probe directed against the *Sal*I repeats^24^. Slides were mounted in 2 µg/ml DAPI diluted in Vectashield (Vectorlabs), imaged on an Axioplan 2 microscope (Carl Zeiss) and acquired with a mono cool view CCD camera. Z-stacks at 100 nm intervals were recorded, deconvolved (AutoQuantX software), slice aligned and Z-projected (HeliconFocus software). RAD51 foci were quantified by manually counting co-localizing signals with the DAPI only. Co-localization with the 45S rDNA probe was scored if the RAD51 focus overlapped by at least 50 % with the labeled probe. Global RAD51 detection was performed as described.

### Protein localization analysis by confocal laser scanning microscopy

Anthers expressing the respective fluorescence reporter construct were dissected, transferred onto a slide with a drop of water and sealed with a cover slip. Images were acquired by using a Leica TCS SP8 inverted confocal microscope or a Zeiss LSM 880 upright microscope, immediately. The fluorescent protein mTurquoise was excited at λ 458 nm and detected at λ 460–510 nm, GFP was excited at 488 nm and detected at 495–560 nm and TagRFP was excited at 561 nm and detected at 570–650 nm.

### STED microscopy

The STED slides were prepared as described for the Immuno-FISH with some minor adjustments. The secondary antibodies used for STED imaging were anti-rat STAR-635P (Abberior) and anti-rabbit STAR-Orange (Abberior). The slides were mounted in Pro-Long Glass antifade (Thermofisher) mounting medium. The super resolution images were acquired with a STED-facility line imaging with a 561 and 640 nm excitation laser with a 775 nm depletion laser.

### RAD21.2 accumulation analysis

To analyze the chromatic features of RAD21.2 accumulations, we performed confocal microscope analysis of meiocytes expressing *PRO_ASK1_GFP:RAD21.2* and *PRO_H2A.W.6_:H2A.W.6:RFP* at pachytene. For 20 meiocytes, 3 areas with no accumulation and 3 areas with accumulations of RAD21.2 were determined. The fluorescence intensity was measured by plot profile in Fiji. For the accumulation evaluation, the maximum intensity of RAD21.2 fluorescence in each of 3 areas was averaged, and relative intensity was calculated as the ratio of the averaged intensity in the RAD21.2 accumulated area to the relative intensity of the non-accumulated area.

### Live cell imaging

Live cell imaging of flower buds was performed according to Pursicki et al.^14^. In brief, a single flower bud was dissected and the stem was embedded into Arabidopsis Apex Culture Medium (APCM) in a petri dish. The sepal was removed to expose two anthers that were covered by a drop of APCM with 2% w/v agarose and the petri dish was filled with autoclaved water and placed under a W-plan Apochromat 40X/1.0 DIC objective. The Zeiss LSM 880 upright confocal microscope and the ZEN 2.3 SP1 software (Carl Zeiss) were used for the acquisition of time lapses. For the analysis of the WAPL dependent removal of RAD21.2, a series of Z-stacks (7 planes, 28 µm distance) were acquired at 15 min time intervals. For the analysis of the RAD21.2 dynamics from premeiosis to pachytene, a series of Z-stacks (10 planes, 45 µm) at 15 min time intervals were acquired.

### Image processing

The time lapses were converted to sequential images and a focal plane was selected for each time point using the function “Review Multi Dimensional Data” of the software Metamorph, version 7.8. Sample drift was corrected by using the Stack Reg plugin of Fiji (version 1.52p)^31^.

For the calculation of the relative intensity of RAD21.2 over the time, time lapses were acquired from leptotene to metaphase I that was denoted as 0 h. We measured the fluorescence intensity of nuclei cross sections from 9-20 meiocytes by using the image processing software Fiji and background fluorescence was subtracted. From the calculated intensity the background intensity was subtracted. The highest measured intensity was marked as 100% and used as reference for the calculation of the RAD21.2 relative intensity for every time point. Representative movies are shown in the Movie S3 (for the wild-type) and Movie S4 (for *wapl1wapl2*).

### Yeast two-hybrid assay

The *SMC1* and *SCC3* constructs were generated as described previously^19^. To generate the *RAD21.2* construct, the coding sequence was amplified by PCR with primers flanking *NdeI* and *NhoI* restriction sites and was subcloned into the *pGADT7* vector by using the T4 Ligase. To generate the *REC8* construct, the coding sequence was amplified by PCR with primers flanked by *attB* sites and subcloned into the *pDONR221* vector by BP clonase reaction. The resulting construct was integrated into the *pGADT7-GW* vector by gateway LR reaction. Primers used for generating the constructs are listed in Supplementary Table S2. The yeast two-hybrid assays were performed according to the Matchmarker Gold Yeast two-hybrid system manual from Clontech. Different variations of the constructs were co-transformed by the polyethylene glycol/ lithium acetate method into the *AH109 yeast* strain and selected on SD/-Leu-Trp plates. The interactions were tested on SD/-Leu-Trp-His plates.

### Plant material collection for protein extraction

2 week old seedlings expressing *PRO_35S_:GFP* or *PRO_ASK1_:GFP:RAD21.2* were grown on ½ MS plates. Around 0.1 g seedlings was collected in a precooled tube and immediately frozen in liquid nitrogen.

### Protein Sample preparation and LC-MS/MS data acquisition

Plant material was ground to a fine powder and covered by the extraction buffer (50 mM Tris pH 7.5, 150 mM NaCl, 10 Glycerol, 2mM EDTA, 5mM DTT, 1% Triton X-100, 10µl/ml plant protease inhibitor (Sigma #P9599)). The extraction was performed for 1 hour on ice with mixing the solution in between. The solution was centrifuged for 30 min at 4°C. The supernatant was collected in a new tube and the centrifugation step was repeated until no pellet was left. For the enrichment, 50 µl of GFP-Trap Magnetic beads (Chromotek) were equilibrated with ice cold wash buffer (50 mM Tris pH 7.5, 150 mM NaCl, 10 Glycerol, 2mM EDETA) according to the manual. Total protein and magnetic beads were mixed and incubated overnight at 4°C on a rolling wheel. The followed wash steps were performed according to the manual and magnetic beads were frozen at -20°C until on-bead digestion was performed. For the on-bead digestion, dry beads were re-dissolved in 25 µL digestion buffer 1 (50 mM Tris, pH 7.5, 2M urea, 1mM DTT, 5 ng/µL trypsin) and incubated for 30 min at 30 °C in a Thermomixer with 400 rpm. Next, beads were pelleted and the supernatant was transferred to a fresh tube. Digestion buffer 2 (50 mM Tris, pH 7.5, 2M urea, 5 mM CAA) was added to the beads, after mixing the beads were pelleted, the supernatant was collected and combined with the previous one. The combined supernatants were then incubated o/n at 32 °C in a Thermomixer with 400 rpm; samples were protected from light during incubation. The digestion was stopped by adding 1 µL TFA and desalted with C18 Empore disk membranes according to the StageTip protocol^32^. Dried peptides were re-dissolved in 2% ACN, 0.1% TFA (10 µL) for analysis and diluted to 0.2 µg/µL . Samples were analyzed using an EASY-nLC 1000 (Thermo Fisher) coupled to a Q Exactive mass spectrometer (Thermo Fisher). Peptides were separated on 16 cm frit-less silica emitters (New Objective, 0.75 µm inner diameter), packed in-house with reversed-phase ReproSil-Pur C18 AQ 1.9 µm resin (Dr. Maisch). Peptides were loaded on the column and eluted for 115 min using a segmented linear gradient of 5% to 95% solvent B (0 min: 5%B; 0-5 min -> 5%B; 5-65 min -> 20%B; 65-90 min ->35%B; 90-100 min -> 55%; 100-105 min ->95%, 105-115 min ->95%) (solvent A 0% ACN, 0.1% FA; solvent B 80% ACN, 0.1%FA) at a flow rate of 300 nL/min. Mass spectra were acquired in data-dependent acquisition mode with a TOP15 method. MS spectra were acquired in the Orbitrap analyzer with a mass range of 300–1750 m/z at a resolution of 70,000 FWHM and a target value of 3×10^6^ ions. Precursors were selected with an isolation window of 2.0 m/z (Q Exactive). HCD fragmentation was performed at a normalized collision energy of 25. MS/MS spectra were acquired with a target value of 10^5^ ions at a resolution of 17,500 FWHM, a maximum injection time (max.) of 120 ms and a fixed first mass of m/z 100. Peptides with a charge of +1, greater than 6, or with unassigned charge state were excluded from fragmentation for MS2, dynamic exclusion for 30s prevented repeated selection of precursors.

### Data analysis

Raw data were processed using MaxQuant software (version 1.5.7.4) with label-free quantification (LFQ) and iBAQ enabled^33^. MS/MS spectra were searched by the Andromeda search engine against a combined database containing the sequences from *A. thaliana* (TAIR10_pep_20101214) and sequences of 248 common contaminant proteins and decoy sequences^34^. Trypsin specificity was required and a maximum of two missed cleavages allowed. Minimal peptide length was set to seven amino acids. Carbamidomethylation of cysteine residues was set as fixed, oxidation of methionine and protein N-terminal acetylation as variable modifications. Peptide-spectrum-matches and proteins were retained if they were below a false discovery rate of 1%. Statistical analysis of the MaxLFQ values was carried out using Perseus (version 1.5.8.5). Quantified proteins were filtered for reverse hits and hits “identified by site” and MaxLFQ values were log2 transformed. After grouping samples by condition only those proteins were retained for the subsequent analysis that had two valid values in one of the conditions. Missing values were imputed from a normal distribution (1.8 downshift, separately for each column). Volcano plots were generated in Perseus using an FDR of 6% and an *S0*=1. Perseus output was exported and further processed using Excel.

### qRT-PCR

Expression analysis of *RAD21.2* in seedlings and flower buds was performed by qRT-PCR. Plant material was collected and grinded to fine powder. RNA extraction was performed according to the manual of the RNeasy Plant Mini kit (Qiagen). A DNase treatment was added before the first washing step. Finally, the RNA concentration was determined and 1 µg RNA was used for the cDNA synthesis according to the Transcriptor First Strand cDNA Synthesis kit (Roche). The expression of the following genes *FTSH7* (AT3G47060), *COX11* (AT1G02410) and AT2G41960 was used as reference. The expression of each gene was analyzed using the primers listed in Supplementary Table S. The qRT-PCR was performed using the Light Cycler 480 SYBR Green I Master (Roche) in triplicates. Following conditions were used; pre-incubation: 95°C 5 min, amplification: 95°C 10 s, 58°C 10 s, 72°C 10 s; 45 cycles. The experiment was performed in a Light Cycler 480 System (Roche).

### Quantitative PCR

4-week-old leaves of T2 *RAD21.2 RNAi* #1 were collected and grinded to fine powder. DNA was extracted by using the DNeasy Plant Pro kit (Qiagen). The qPCR was performed in triplicates and 1.5 ng DNA was used. To quantify the relative *18S* gene number primers, previously described, were used^21^. To calculate the relative 18S quantity the *HXK1* (AT4G29130) and *UEV1C* (AT2G36060) genes were used for normalization. Following conditions were used; pre-incubation: 95°C 7 min, amplification: 95°C 30 s, 56°C 30 s, 72°C 30 s; 40 cycles. The experiment was performed in a Light Cycler 480 System (Roche).

For detecting the copy number variation within the same individual, rosette leaves of different sizes, representing different ages, were collected from bottom to top as follows t0 = 1cm leaves, t1 = 1.5 cm, t2 = 2.5 cm, t3 = 3.5 cm. Four leaves per time point per individual were taken for the analysis.

### Statistical analysis

Student’s t-test (two-tailed) was used to evaluate the significance of the difference between two groups. For the analysis of variance, two samples F-test was performed. The numbers of samples are indicated in the figure legend. The strength of significance is presented by the p-values. *,P< 0.05; **,P< 0.01; and ***,P< 0.001. Unpaired, two-tailed Mann-Whitney tests were performed, since D’Agostino Pearson omnibus K2 normality testing revealed that most data were not sampled from a Gaussian population, and nonparametric tests were therefore required.

## Tables S1 to S3

**Table S1: Proteins identified by IP_RAD21.2 vs. GFP**

**Table S2: Oligonucleotides used in this study**

## Figs. S1 to S7

**Fig. S1.**
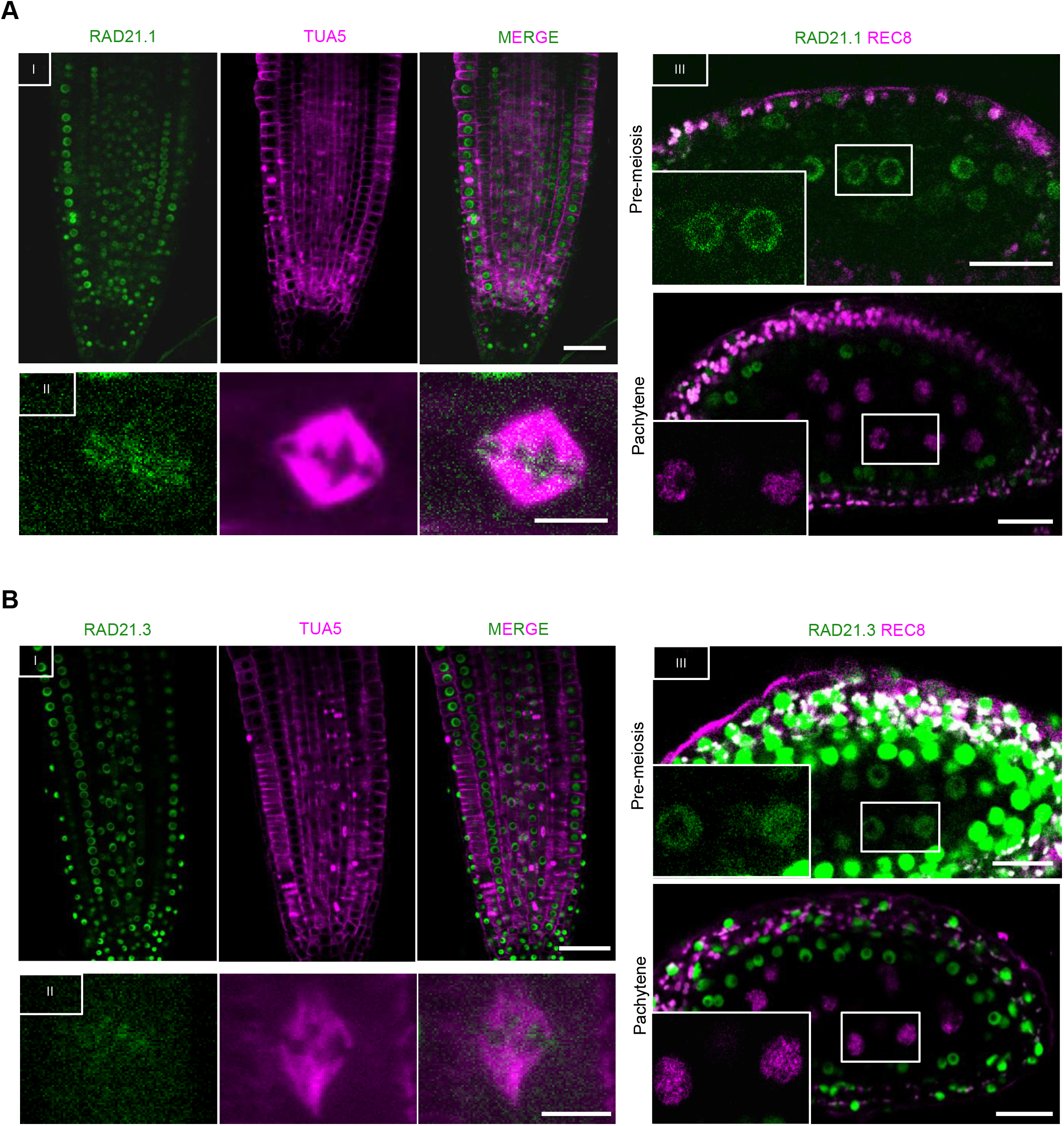
Mitotic and meiotic localization of RAD21.1 and RAD21.3: Confocal laser scanning micrographs of Arabidopsis root tips (I, II) and anthers (III) expressing *PRO_RPS5_:RFP:TUA5* together with *PRO_RAD21.1_:RAD21.1:GFP* (**A**) and *PRO_RAD21.3_:GFP:RAD21.3* (**B**). (I) Depicts an overview of the root tip. Scale bar: 50 µm. (II) Close-up showing the localization pattern of RAD21 fusion proteins on chromosomes in the metaphase plane. Scale bar: 5 µm. (III) Upper row: RAD21.1 and RAD21.3 but not REC8 are present in all cells prior to meiosis (close-up highlighted in box). Lower row: next to REC8, neither RAD21.1 nor RAD21.3 accumulate in meiosis (here pachytene stage) but are present in the surrounding somatic tissue. Scale bar: 20 µm.

**Fig. S2.**
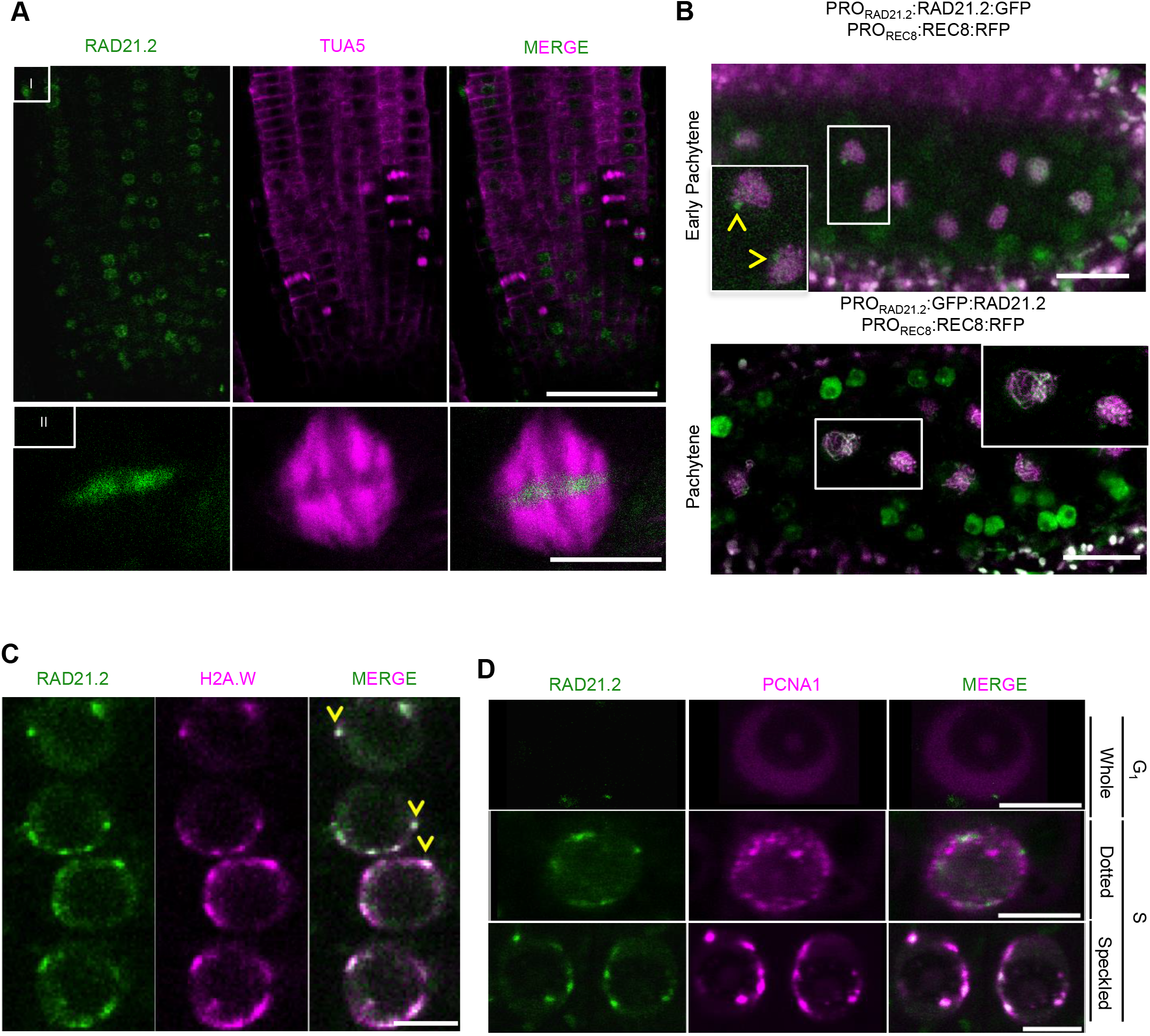
RAD21.2 localization in Arabidopsis root cells overlaps with H2A.W.6: **A,** Confocal laser scanning micrographs of Arabidopsis root tips (I) expressing *PRO_RPS5_:RFP:TUA5* together with *PRO_RAD21.2_:GFP:RAD21.2*. (I) Overview of the root tip. Scale bar: 50 µm. (II) Close-up showing the localization pattern of RAD21 fusion proteins on chromosomes in the metaphase plane. Scale bar: 5 µm. **B**, Upper panel: Confocal laser scanning micrograph of an anther expressing PRO_RAD21.2_:RAD21.2:GFP (green) and PRO_REC8_:REC8:GFP (magenta) at early pachytene stage. Lower panel: Confocal laser scanning micrograph of an anther expressing PRO_RAD21.2_:GFP:RAD21.2 (green) and PRO_REC8_:REC8:GFP (magenta) at pachytene stage. Arrowheads show RAD21.2 clusters. Scale bar: 20 µm. **C,** Confocal laser scanning micrographs of Arabidopsis root tip cells expressing *PRO_ASK1_:GFP:RAD21.2* (green) and *PRO_H2A.W.6_:H2A.W.6:RFP* (magenta) in root cells. Overlapping regions are marked with arrowheads. Scale bar: 5 µm. **D,** Confocal laser scanning micrographs of Arabidopsis root tip cells expressing *PRO_RAD21.2_:GFP:RAD21.2* (green) and *PRO_PCNA1_:PCNA1:RFP* (magenta) depicting cell cycle-dependent dynamics of RAD21.2 in root cells. Scale bar: 5 µm.

**Fig. S3.**
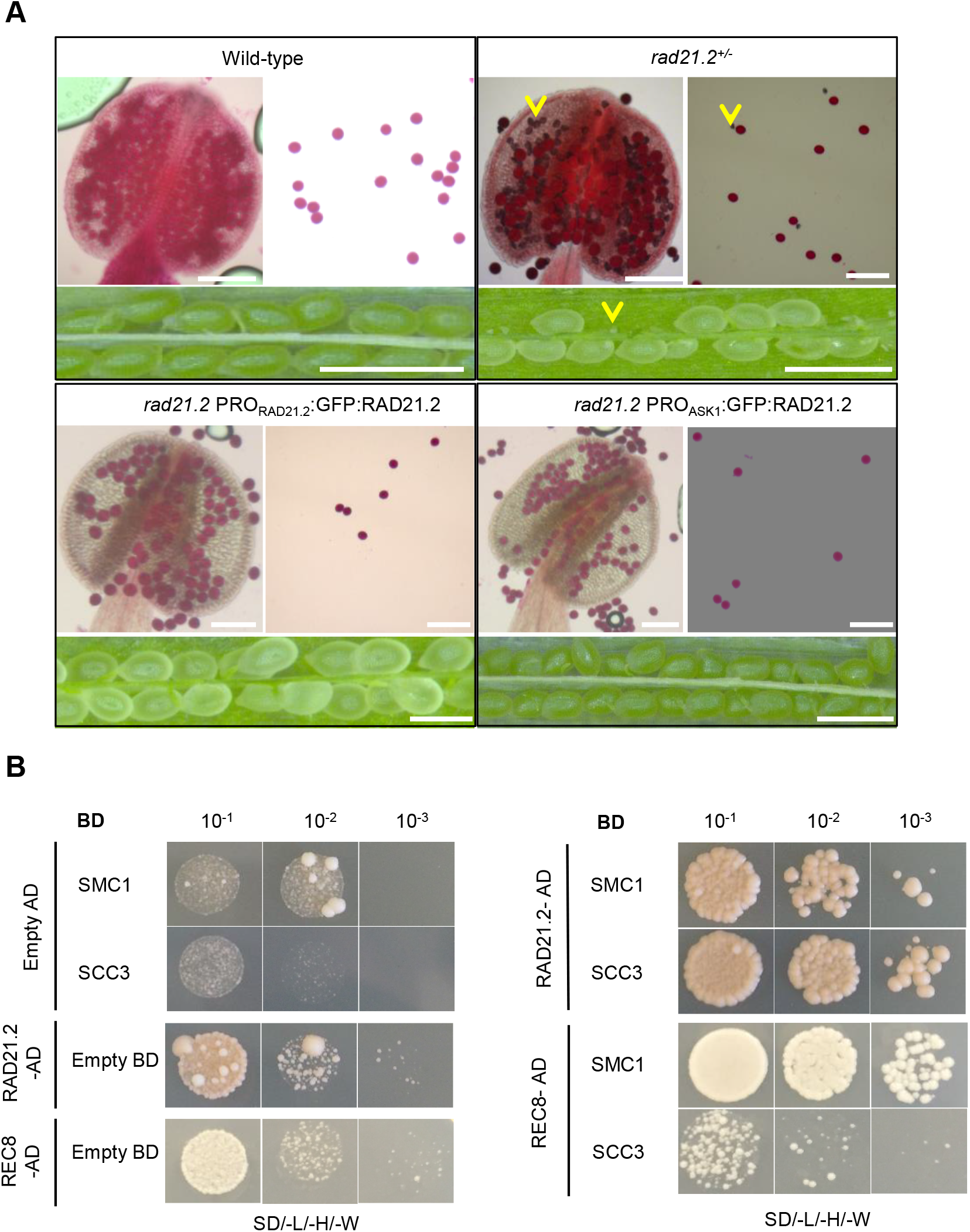
GFP:RAD21.2 reporters complement the *rad21.2* phenotype and show a specific pre- and post-meiotic localization pattern: **A**, *RAD21.2* reporter complementation assays. Peterson staining of mature pollen. Aborted pollen is identifiable by blue color and shrunken appearance (arrowhead). Aborted seeds are highlighted by arrowheads. Phenotypes of the wild-type, *rad21.2* heterozygous mutants, and plants carrying either of the reporter constructs *PRO_RAD21.2_:GFP:RAD21.2* and *PRO_ASK1_:GFP:RAD21.2*. Heterozygous mutants for *RAD21.2* show a 40% pollen and 50% seed viability reduction. Scale bar for seed analysis: 1000 µm; scale bar for pollen analysis: 100 µm. **B,** Yeast two-hybrid interaction assay of RAD21.2 and REC8 with the core cohesin components SMC1 and SCC3. The left panel shows the autoactivation tests for all analyzed constructs. The right panel shows the results of the interaction analyses. Different dilutions of yeast (10^-1^/10^-2^/10^-3^) were spotted on SD plates lacking leucine, tryptophan and histidine (-L/-W/-H) to test for interaction strength.

**Fig. S4.**
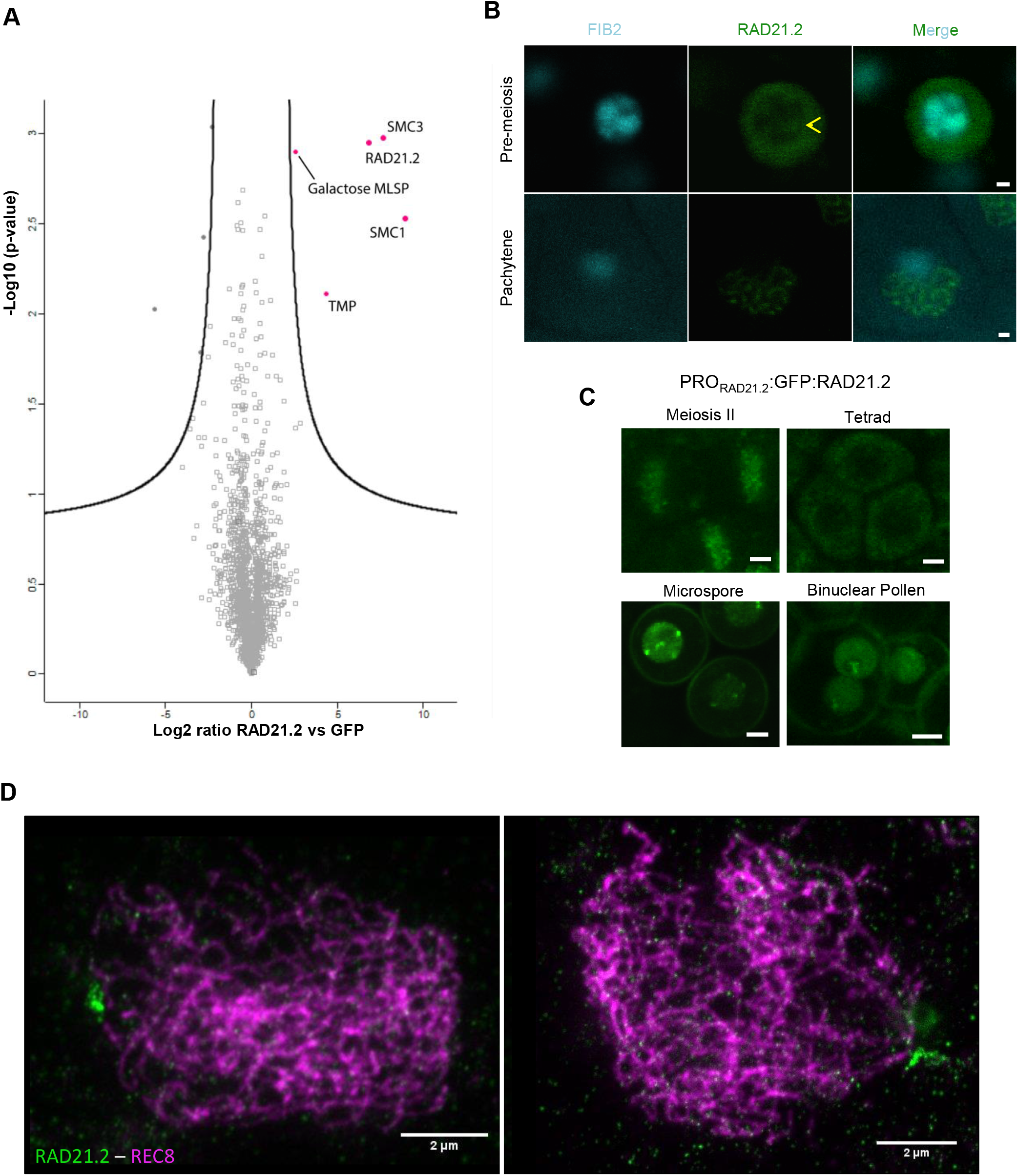
RAD21.2 interacts with core cohesin components and co-precipitates with proteins associated with heterochromatin: **A,** Volcano plot of proteins identified by mass spectrometry. The x-axis depicts the fold change value, and the y-axis shows the significance by the –log10 (p-value). Proteins were extracted from seedlings. Interaction partners that are significantly enriched in the RAD21.2 IP sample are shown in magenta. Other identified proteins are shown in grey (Table S1). **B**, Confocal laser scanning micrographs of cells expressing PRO_ASK1_:GFP:RAD21.2 (green) and PRO_FIB2_:FIB2:TFP (cyan) in pre-meiosis indicating a nuclear localization of RAD21.2 with nucleolar protruding regions (arrowhead) at pre-meiosis (upper panel) and non-protruding regions at pachytene stage (lower panel). Scale bar: 1 µm. **C**, Confocal laser scanning micrographs of cells expressing PRO_RAD21.2_:GFP:RAD21.2 (green) at the end of meiosis II, tetrad, microspore and bicellular pollen stages. Scale bar: 1 µm. **D**, Two representative images of Lipsol spread nuclei imaged in super-resolution using STED microscopy. REC8 is shown in magenta and RAD21.2 in green. Nuclei are at zygotene stage. Scale bar: 2 µm.

**Fig. S5.**
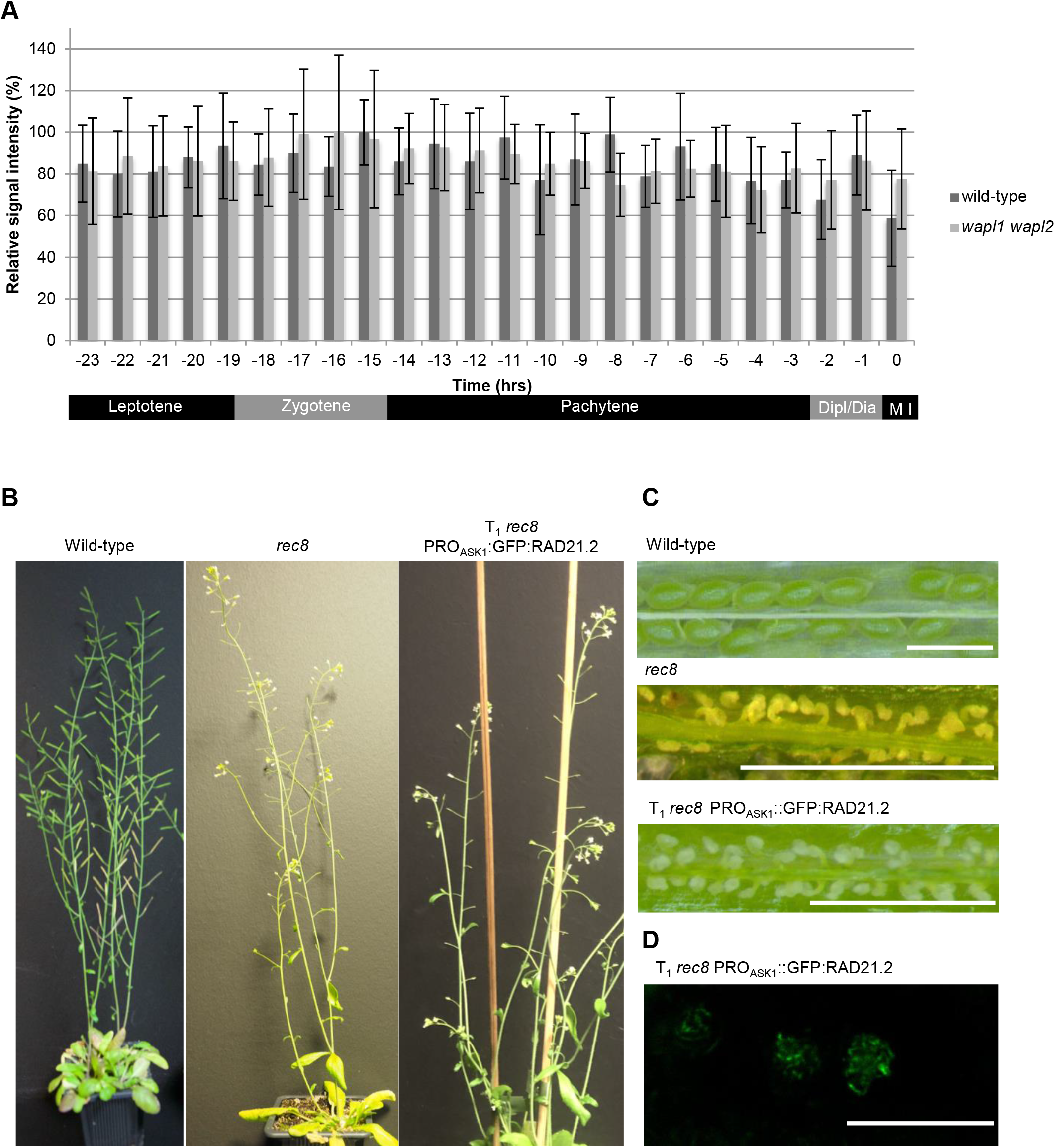
RAD21.2 is not regulated by the WAPL-dependent prophase I pathway and RAD21.2 localization is REC8-independent: **A,** Quantification of the relative PRO_ASK1_:GFP:RAD21.2 accumulation levels in wild-type (dark grey) versus *wapl1 wapl2* (light grey) meiocytes over time. At least 9 meiocytes were analyzed per genotype, error bars depict the standard error. Meiotic stages are indicated below (see Movies S3 and S4). **B**, Expression of *PRO_ASK1_:GFP:RAD21.2* does not complement the sterility of *rec8* mutants as seen by the short (**B**) and empty (**C**) siliques. Scale bar: 1000 µm. Description for panel C is missing. **D**, Confocal laser scanning micrograph of an anther expressing *PRO_ASK1_:GFP:RAD21.2* in *rec8* shows the typical chromosomal localization pattern of GFP:RAD21.2 at pachytene; see Fig. 1 for comparison.

**Fig. S6.**
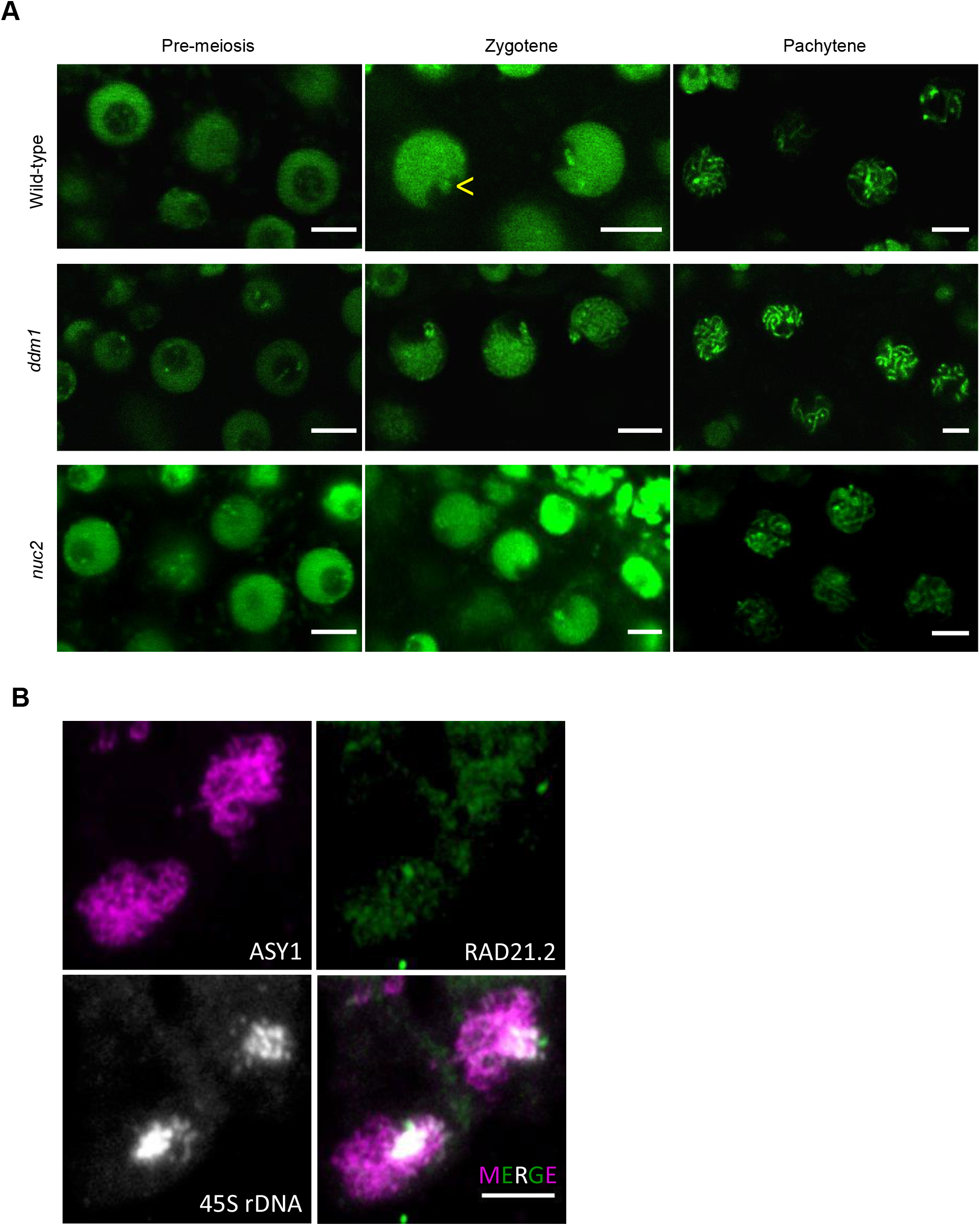
RAD21.2 localization is not compromised in *ddm1* and *nuc2* mutant meiocytes: **A,** Confocal laser scanning micrographs of anthers expressing *PRO_ASK1_:GFP:RAD21.2* in *ddm1* and *nuc2* mutants showing the typical chromosomal localization pattern of GFP:RAD21.2 (green) from pre-meiosis to pachytene compared to wild-type. B, Immuno-FISH analysis of wild-type pollen mother cells at zygotene. The axis has been stained with anti-ASY1 (magenta) for staging and the RAD21.2 by anti-RAD21.2 (green). The 45S rDNA has been visualized with a specific FISH probe (white). Scale bar: 5 µm.

**Fig. S7.**
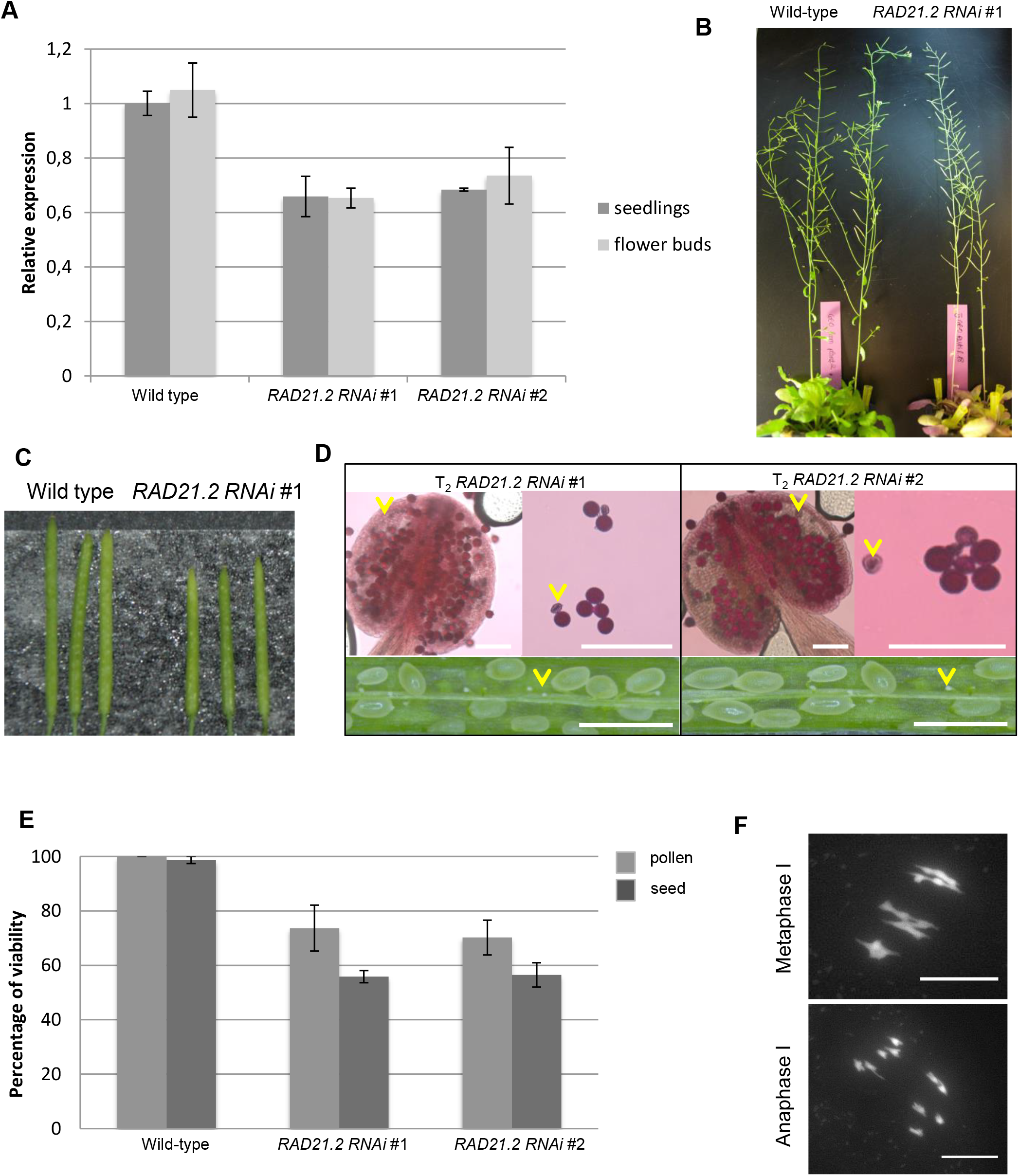
Knockdown of *RAD21.2* results in reduced fertility: **A,** Normalized relative expression levels of *RAD21.2* in seedlings (dark grey) and in flower buds (light grey) in the *RAD21.2 RNAi* lines #1 and #2 compared to the wild type. **B**, **C** *RAD21.2* RNAi plants (line #1) have somatically visible defects but have shorter siliques than the wild type. **D**, *RAD21.2 RNAi* plants (line #1 left, and line #2 right) have dead pollen (examples marked by arrowheads) and aborted and unfertilized ovules (examples are marked by arrowheads). Scale bar for silique analysis: 1000 µm, scale bar for pollen analysis: 100 µm. **E**, Quantification of the fertility defects in *RAD21.2 RNAi* plants. **F**, F1 cross between *RAD21.2* RNAi #1 and Col-0 show no severe paring defects indicating that no translocation events have occurred while selecting for the *RAD21.2* RNAi line.

**Fig. S8.**
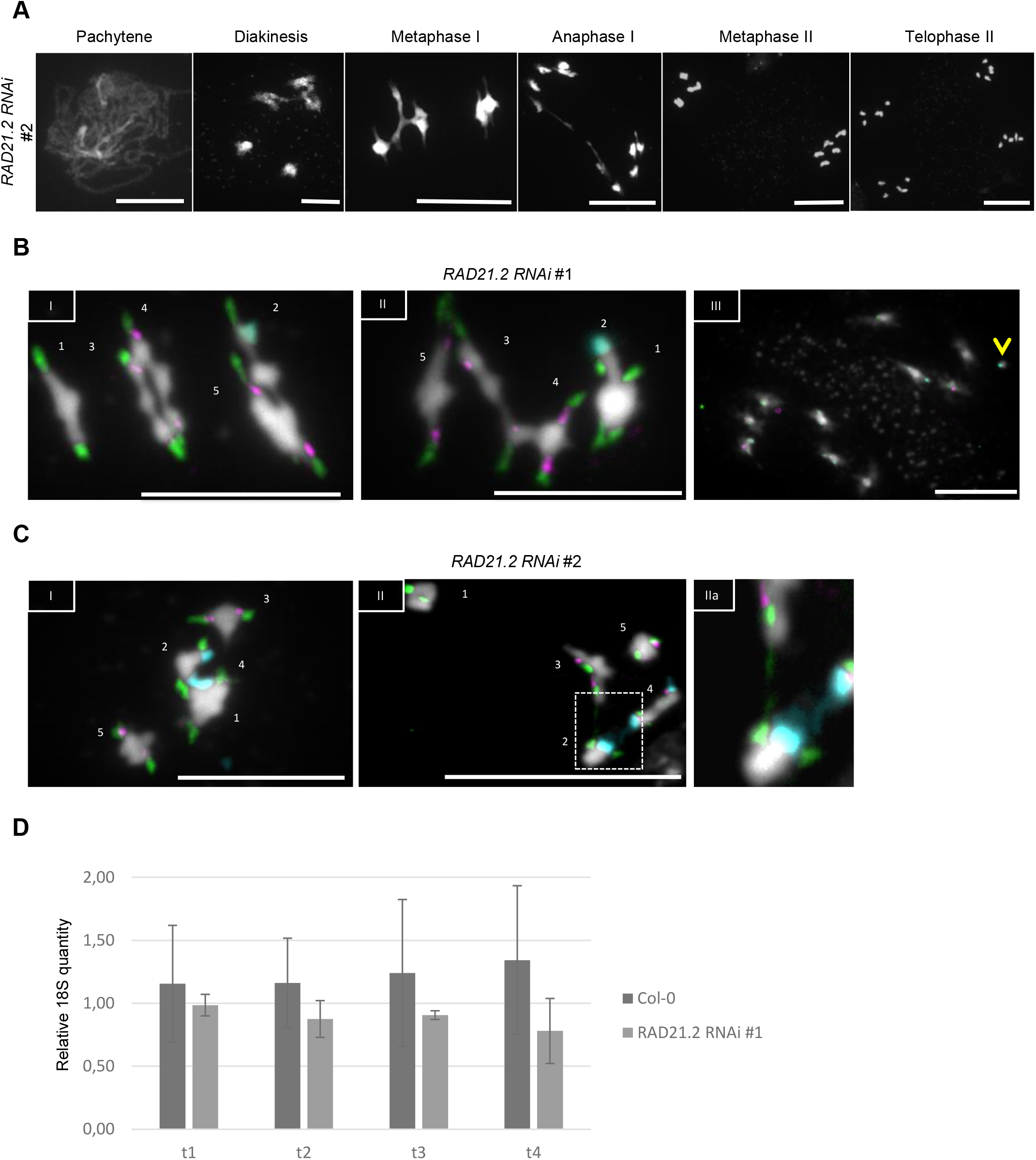
*RAD21.2 RNAi* plants have severe chromosome rearrangements: **A,** Chromosome spread analysis of pollen mother cells of the *RAD21.2 RNAi* line #2. Scale bar: 20 µm. **B,** FISH analysis of metaphase I (I/II) and prophase II (III) of pollen mother cells from *RAD21.2 RNAi* line #1. DNA was visualized by DAPI (grey), probes against the 45S rDNA (cyan), 5S rDNA (red) and CEN (green) regions were used to identify chromosomes. Several arrangements of non-homologous chromosome were found, e.g. chromosomes 3 and 4 (I) and chromosome 5 with 3 and 3 with 4 (II). An arrowhead indicates a 45S chromosome fragment (III). Scale bar: 10 µm. **C,** FISH analysis of metaphase I (I/II) of pollen mother cells from *RAD21.2 RNAi* line #2. DNA was visualized by DAPI (grey), probes against the 45S rDNA (cyan), 5S rDNA (red) and CEN (green) regions were used to identify chromosomes. Interconnections of chromosomes 2 and 4 were visible (I). Panel (IIa) depicts the magnification of the marked area in (II) highlighting connections between chromosomes 2 and 4 via the 45S region and between chromosomes II and III via the centromeric region. Scale bar: 10 µm. **D**, Graph depicting the 18S gene copy number in leaves of different sizes of Col-0 and *RAD21.2 RNAi* line #1. The results show that there is loss of 18S copy number occurring within the same individual, indicating there is no somatic loss of the rDNA.

## Movies S1 to S4

**Movie S1. Dynamics of RAD21.1 in mitotically dividing root cells**: Live cell imaging of root tips expressing PRO_RPS5_:RFP:TUA5 (magenta) together with PRO_RAD21.1_:RAD21.1:GFP (green). A typical cohesin localization pattern was observed, characterized by chromatin association and disappearance at anaphase onset. The fluorescence signal of PRO_RAD21.1_:RAD21.1:GFP reappears in the nuclei of the daughter cells after mitosis. Time interval of image acquisition is 20 s.

**Movie S2. Pre-meiotic RAD21.2 dynamics**: Live cell imaging of anthers expressing *PRO_ASK1_:GFP:RAD21.2*. RAD21.2 (grey) accumulates in small foci in the nuclei of pre-meiotic cells and the surrounding tapetum cells. Time interval of image acquisition is 15 min.

**Movie S3. RAD21.2 dynamics from leptotene to anaphase I in the wild-type**: Live cell imaging of wild-type flower buds expressing PRO_ASK1_:GFP:RAD21.2. RAD21.2 (grey) is enriched at distinct chromosome regions. Time interval of image acquisition is 15 min.

**Movie S4. RAD21.2 dynamics from leptotene to anaphase I in *wapl1 wapl2* double mutants**: Live cell imaging of *wapl1 wapl2* flower buds expressing PRO_ASK1_:GFP:RAD21.2. Loss of WAPL function does not affect the localization and abundance pattern of RAD21.2 (grey). Time interval of image acquisition is 15 min.

## References

1. Nasmyth, K. & Haering, C. H. Cohesin: its roles and mechanisms. Annu Rev Genet 43, 525–558 (2009).

2. Klein, F. et al. A central role for cohesins in sister chromatid cohesion, formation of axial elements, and recombination during yeast meiosis. Cell 98, 91–103 (1999).

3. Watanabe, Y. & Nurse, P. Cohesin Rec8 is required for reductional chromosome segregation at meiosis. Nature 400, 461–464 (1999).

4. Bhatt, A. M. et al. The DIF1 gene of Arabidopsis is required for meiotic chromosome segregation and belongs to the REC8/RAD21 cohesin gene family. The Plant Journal 19, 463–472 (1999).

5. Severson, A. F., Ling, L., van Zuylen, V. & Meyer, B. J. The axial element protein HTP-3 promotes cohesin loading and meiotic axis assembly in C. elegans to implement the meiotic program of chromosome segregation. Genes & development 23, 1763–1778 (2009).

6. Severson, A. F. & Meyer, B. J. Divergent kleisin subunits of cohesin specify mechanisms to tether and release meiotic chromosomes. Elife 3, e03467 (2014).

7. Herrán, Y. et al. The cohesin subunit RAD21L functions in meiotic synapsis and exhibits sexual dimorphism in fertility. The EMBO journal 30, 3091–3105 (2011).

8. Ishiguro, K., Kim, J., Fujiyama-Nakamura, S., Kato, S. & Watanabe, Y. A new meiosis-specific cohesin complex implicated in the cohesin code for homologous pairing. EMBO Rep 12, 267–275 (2011).

9. Lee, J. & Hirano, T. RAD21L, a novel cohesin subunit implicated in linking homologous chromosomes in mammalian meiosis. J Cell Biol 192, 263–276 (2011).

10. da Costa-Nunes, J. A. et al. Characterization of the three Arabidopsis thaliana RAD21 cohesins reveals differential responses to ionizing radiation. J Exp Bot 57, 971–983 (2006).

11. Jiang, L., Xia, M., Strittmatter, L. I. & Makaroff, C. A. The Arabidopsis cohesin protein SYN3 localizes to the nucleolus and is essential for gametogenesis. The Plant Journal 50, 1020–1034 (2007).

12. Yuan, L., Yang, X., Ellis, J. L., Fisher, N. M. & Makaroff, C. A. The Arabidopsis SYN3 cohesin protein is important for early meiotic events. Plant J 71, 147–160 (2012).

13. Yuan, L., Yang, X., Auman, D. & Makaroff, C. A. Expression of epitope-tagged SYN3 cohesin proteins can disrupt meiosis in Arabidopsis. J Genet Genomics 41, 153–164 (2014).

14. Prusicki, M. A. et al. Live cell imaging of meiosis in Arabidopsis thaliana. Elife 8, (2019).

15. Pontvianne, F. et al. Identification of nucleolus-associated chromatin domains reveals a role for the nucleolus in 3D organization of the A. thaliana genome. Cell reports 16, 1574–1587 (2016).

16. Ingouff, M. et al. Live-cell analysis of DNA methylation during sexual reproduction in Arabidopsis reveals context and sex-specific dynamics controlled by noncanonical RdDM. Genes Dev 31, 72–83 (2017).

17. Yelagandula, R. et al. The histone variant H2A.W defines heterochromatin and promotes chromatin condensation in Arabidopsis. Cell 158, 98–109 (2014).

18. De, K., Sterle, L., Krueger, L., Yang, X. & Makaroff, C. A. Arabidopsis thaliana WAPL is essential for the prophase removal of cohesin during meiosis. PLoS Genet 10, e1004497 (2014).

19. Yang, C. et al. SWITCH 1/DYAD is a WINGS APART-LIKE antagonist that maintains sister chromatid cohesion in meiosis. Nat Commun 10, 1755 (2019).

20. Concia, L. et al. “Genome-wide analysis of the Arabidopsis replication timing program.” Plant physiology 176.3 (2018).

21. Soppe, Wim JJ, et al. DNA methylation controls histone H3 lysine 9 methylation and heterochromatin assembly in Arabidopsis. The EMBO journal 21.23, 6549–6559(2002).

22. Hirochika, Hirohiko, Hiroyuki Okamoto, and Tetsuji Kakutani. Silencing of retrotransposons in Arabidopsis and reactivation by the ddm1 mutation. The Plant Cell 12.3, 357–368 (2000).

23. Osakabe, Akihisa, et al. The chromatin remodeler DDM1 prevents transposon mobility through deposition of histone variant H2A. W. Nature Cell Biology 23.4, 391–400 (2021).

24. Durut, Nathalie, et al. A duplicated NUCLEOLIN gene with antagonistic activity is required for chromatin organization of silent 45S rDNA in Arabidopsis. The Plant Cell 26.3, 1330–1344 (2014).

25. Mozgová, Iva, Petr Mokroš, and Jiří Fajkus. Dysfunction of chromatin assembly factor 1 induces shortening of telomeres and loss of 45S rDNA in Arabidopsis thaliana. The Plant Cell 22.8, 2768–2780 (2010).

26. Muchová, Veronika, et al. Homology-dependent repair is involved in 45 S rDNA loss in plant CAF-1 mutants. The Plant Journal 81.2, 198–209 (2015).

27. Sims, J., Copenhaver, G. P. & Schlögelhofer, P. Meiotic DNA Repair in the Nucleolus Employs a Nonhomologous End-Joining Mechanism. Plant Cell 31, 2259–2275 (2019).

28. Wellmer, F., Alves-Ferreira, M., Dubois, A., Riechmann, J. L. & Meyerowitz, E. M. Genome-wide analysis of gene expression during early Arabidopsis flower development. PLoS Genet 2, e117 (2006).

29. Komaki, S. & Schnittger, A. The Spindle Assembly Checkpoint in Arabidopsis Is Rapidly Shut Off during Severe Stress. Dev Cell 43, 172–185.e5 (2017).

30. Ross, K. J., Fransz, P. & Jones, G. H. A light microscopic atlas of meiosis in Arabidopsis thaliana. Chromosome Res 4, 507–516 (1996).

31. Schindelin, J., et al. Fiji: an open-source platform for biological-image analysis. Nat Methods 9, 676–682 (2012).

32. Rappsilber, J., Ishihama, Y. & Mann, M. Stop and go extraction tips for matrix-assisted laser desorption/ionization, nanoelectrospray, and LC/MS sample pretreatment in proteomics. Analytical chemistry 75, 663–670 (2003).

33. Cox, J. & Mann, M. MaxQuant enables high peptide identification rates, individualized p.p.b.-range mass accuracies and proteome-wide protein quantification. Nat Biotechnol 26, 1367–1372 (2008).

34. Tyanova, S., Temu, T. & Cox, J. The MaxQuant computational platform for mass spectrometry-based shotgun proteomics. Nat Protoc 11, 2301–2319 (2016).

